# Reverse signaling by Semaphorin-6A regulates cellular aggregation and neuronal morphology

**DOI:** 10.1101/036095

**Authors:** Francesc Perez-Branguli, Yvrick Zagar, Daniel K. Shanley, Isabella A. Graef, Alain Chédotal, Kevin J. Mitchell

**Affiliations:** Smurfit Institute of Genetics and Institute of Neuroscience, Trinity College Dublin, Dublin 2, Ireland; Sorbonne Universités, UPMC Univ Paris 06, INSERM UMR_S968, CNRS_UMR7210, Institut de la Vision, F-750012, Paris, France; Department of Pathology, Stanford University Medical School, Stanford, CA 94305, USA.

**Keywords:** Abl, plexin, reverse, semaphorin, signaling

## Abstract

The transmembrane semaphorin, Sema6A, has important roles in axon guidance, cell migration and neuronal connectivity in multiple regions of the nervous system, mediated by context-dependent interactions with plexin receptors, PlxnA2 and PlxnA4. Here, we demonstrate that Sema6A can also signal cell-autonomously, in two modes, constitutively, or in response to higher-order clustering mediated by either PlxnA2-binding or chemically induced multimerisation. Sema6A activation stimulates recruitment of Abl to the cytoplasmic domain of Sema6A and phosphorylation of this cytoplasmic tyrosine kinase, as well as phosphorylation of additional cytoskeletal regulators. Sema6A reverse signaling affects the surface area and cellular complexity of non-neuronal cells and aggregation and neurite formation of primary neurons *in vitro*. Sema6A also interacts with PlxnA2 in *cis*, which reduces binding by PlxnA2 of Sema6A *in trans* but not vice versa. These experiments reveal the complex nature of Sema6A biochemical functions and the molecular logic of the context-dependent interactions between Sema6A and PlxnA2.

## Introduction

Morphogenesis of the nervous system requires the migration of myriad cell types to their preordained positions, the guidance of growing axons along stereotyped pathways to their final target regions and the pairing of pre- and postsynaptic cellular partners. Semaphorins comprise a large family of proteins with important roles in many of these processes, coordinated with the families of their receptor proteins, neuropilins and plexins [1].

The transmembrane Semaphorin-6 subclass consists of four members, which interact directly with members of the Plexin-A subclass. Of these, Semaphorin-6A (Sema6A) and Semaphorin-6B (Sema6B) form a cognate subgroup with Plexin-A2 (PlxnA2) and Plexin-A4 (PlxnA4). Genetic studies have revealed context-dependent interactions between members of this group, which control processes of cell migration, axon guidance and neuropil organization in various parts of the developing nervous system [1]. Sema6A signals cell-non-autonomously via PlxnA2 on responding cells to initiate a switch in migratory mode in cerebellar granule cells [2,3] and to restrict motor neuron cell bodies from exiting the ventral nerve root in the spinal cord [4,5]. Sema6A signals via PlxnA4, to confine corticospinal projections [6,7] and to establish laminar connectivity in the retina [8,9]. In the developing hippocampus, signals from both Sema6A and Sema6B restrict mossy fibre projections via PlxnA4 [10,11].

In addition to trans interactions across cells, *cis* interactions are also important. In the hippocampus, PlxnA2 co-expression in the target zone antagonizes Sema6A-PlxnA4 signaling and defines a permissive zone for mossy fibre projection and synapse formation [10,11]. *Cis* interactions between Sema6A and PlxnA2 may also be important in controlling dendritic arborization of retinal ganglion cells [12]. Direct binding in *cis* has also been demonstrated between Sema6A and PlxnA4. In sensory neurons, co-expression of Sema6A inhibits the response of PlxnA4 to Sema6A in trans, thus making these neurons insensitive to this cue, unlike sympathetic neurons, which normally express only PlxnA4 [13]. The fact that the four proteins in this cognate group are often co-expressed suggests that such *cis* interactions may contribute significantly to the combinatorial logic of their functions in other contexts.

Signaling may also occur in a bidirectional manner between these proteins. Interactions in the canonical “forward” direction, with Sema6A or Sema6B as the ligand and PlxnA2 and/or PlxnA4 in the responding cells, cannot readily account for all the phenotypes observed in animals with mutations in these genes, many of which are apparent in axons normally expressing Sema6A [14,15] or co-expressing Sema6A and PlxnA2 [12]. Moreover, there are several known examples of related transmembrane semaphorins that can also signal in the “reverse” direction, mediating cell-autonomous responses to exogenous cues.

In Drosophila, Sema1a, the orthologue of the Sema6 family, acts as a receptor for Plexin-A (PlexA) in the guidance of photoreceptor axons [16,17]. In the olfactory system, Sema1a acts as a ligand for PlexA to control targeting of odorant receptor neuron axons [18], but as a receptor for the secreted semaphorins Sema2a and Sema2b to control dendritic targeting of projection neurons [19]. In development of giant fibre connectivity and motor axon projections, Sema1a signals in a bidirectional manner simultaneously [20,21], though the ligand that activates Sema1a reverse signaling in these contexts is unknown. Similarly, in vertebrates, Sema6D has been shown to signal in a bidirectional manner, acting as both ligand and receptor for PlxnA1 during heart development [22].

Information on the downstream signaling pathways is more limited for semaphorin reverse signaling than for forward signaling through plexin and neuropilin receptors [23,24]. In flies, an interaction between the cytoplasmic domain of Sema1a and the cytoskeletal regulator protein Enabled (Ena) is required for reverse signaling [20]. Sema1a signals are also transmitted via regulators of Rho family small GTPases [21] to control axon guidance. Interactions in *cis* with the L1 orthologue neuroglian may modulate Sema1a receptor function [25]. In vertebrates, Sema6D activation leads to recruitment of the cytoplasmic tyrosine kinase Abl to the Sema6D cytoplasmic domain and phosphorylation of Ena [22]. Sema6A has been shown to bind the Ena paralogue, Evl [26] and computational analysis strongly predicts an Abl-binding site in the Sema6A cytoplasmic domain [27]. Both the Abl and Ena/Evl-binding site sequences are highly conserved between Sema6A and Sema6D. Finally, Sema6B has been reported to bind the cytoplasmic tyrosine kinase Src [28].

Crystal structures have illustrated the mechanism whereby binding of Sema6A activates forward signaling through PlxnA2 [29,30]. Sema6A dimers independently bind two PlxnA2 monomers, followed by higher-order clustering in the membrane, possibly mediated by *cis* dimerization of PlxnA2 through the membrane-proximal IPT domains or intracellular regions [31] or trimerization in a complex with a RhoGTPase [32,33]. This clustering activates PlxnA2 intracellular signaling, but whether it also activates signaling in the reverse direction, through the Sema6A cytoplasmic domain, remains unknown.

Here, we show that Sema6A is capable of reverse signaling, both constitutively and in response to either PlxnA2-binding or chemically induced multimerisation. Activation of Sema6A stimulates phosphorylation of numerous downstream signaling proteins, including Abl kinase, which is recruited to the Sema6A cytoplasmic domain on ligand binding. Sema6A reverse signaling affects the shape of non-neuronal cells and aggregation and neurite formation of neurons *in vitro*. Sema6A also interacts with PlxnA2 in *cis*, which reduces binding by PlxnA2 of Sema6A in *trans* but not vice versa. These experiments reveal the complex nature of Sema6A biochemical functions and begin to dissect the molecular logic of the context-dependent interactions between Sema6A and PlxnA2.

## Results

### Sema6A signals cell-autonomously in NIH3T3 cells and confers responsiveness to PlxnA2

To address the potential cell-autonomous functions of Sema6A we performed transfection of Sema6A in NIH3T3 cells. Cells expressing the full-length Sema6A (Sema6A-FL) showed the constitutive appearance of multiple membrane filopodia around the cell perimeter (Figure 1a and b). These results indicate the Sema6A homodimer has a constitutive cell-autonomous activity in mouse cells under these conditions. Incubation of 60–100nM of PlxnA2 extracellular domain (PlxnA2-EC) had no effect on untransfected cells but on cells expressing Sema6A-FL promoted internalization of Sema6A in vesicle-like structures, together with a reduction of membrane protrusions and cellular contraction, along with collapse of F-actin (Figure 1b, e and g; Figure S1a-f). PlxnA2-EC was able to provoke this cell collapse in cells expressing Sema6A-FL but not cells expressing Sema6B-FL or Sema6D-FL (Figure S1m). Lysine 393 is in the PlxnA2-binding interface of Sema6A and mutation of this residue to aspartic acid (K393D) abrogates PlxnA2 binding [29,30]. Expression of Sema6A carrying the K393D mutation (Sema6A-K393D) still led to cellular expansion and filopodia formation but did not confer responsiveness to PlxnA2 (Figure 1c, f and g; Figure S1g-l).

**Figure 1.**
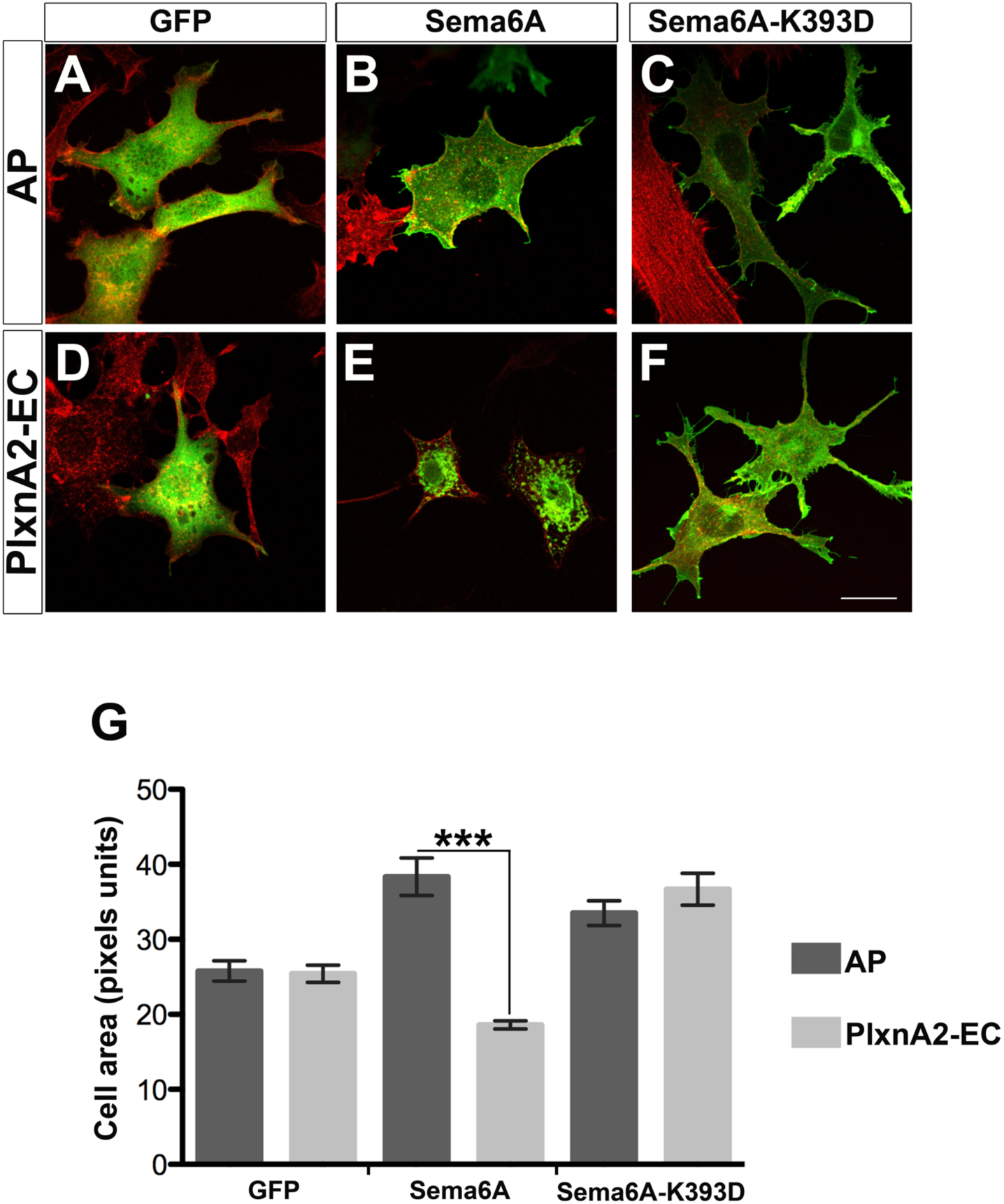
Sema6A exerts constitutive and PlxnA2-dependent cell-autonomous functions. **(a-f)** NIH3T3 cells expressing GFP alone **(a and b)**, myc-Sema6A-FL (Sema6A; **b and e)**, or myc-Sema6A-K393D (Sema6A-K393D; **c and f)** were treated with purified AP-Fc (AP; **a-c)** or PlxnA2-EC-Fc (PlxnA2-EC; **d-f)**. Scale bar = 20 μm. **(g)** Graph represents the cell area in cells transfected with GFP, Sema6A or Sema6A-K393D and treated with AP or PlxnA2-EC; *n*=100–200 cells per experimental condition. Data are expressed as mean ± s.e.m; \*\*\**p*< 0.001; Student’s *t*-test.

### PlxnA2-induced Sema6A reverse signaling regulates aggregation of cerebellar granule cells

Forward signaling from Sema6A to PlxnA2 plays a crucial role in the development of the cerebellum, controlling the radial migration of the granule cell neurons from the external granule cell layer (EGL) [2,3]. We used explants of cerebellar granule cells to explore the consequences of signaling in the reverse direction. We previously showed that culturing EGL explants on NIH3T3 cells expressing Sema6A causes a reduction in the number of migrating neurons (Figure 2b) in a PlxnA2-dependent manner (Figure 2f) [29]. In contrast, explants cultured on PlxnA2-expressing NIH3T3 cells revealed a high increase in the number of cells outside the explant and apparent disaggregation of the explant core (Figure 2c). Explants showed no response to cells expressing the A396E mutant form of PlxnA2 (PlxnA2-A396E; Figure 2d), which is incapable of binding Sema6A [3,29,30]. The response to PlxnA2 was dependent on expression of Sema6A in the EGL neurons, as explants from *Sema6A^/-^* animals showed no effect (Figure 2k). Mutation of Sema6A in the granule cells had no effect on response to Sema6A in *trans* (Figure 2j), nor did mutation of PlxnA2 affect the response to PlxnA2 in *trans* (Figure 2g). These results are quantified in Figure 2m.

**Figure 2.**
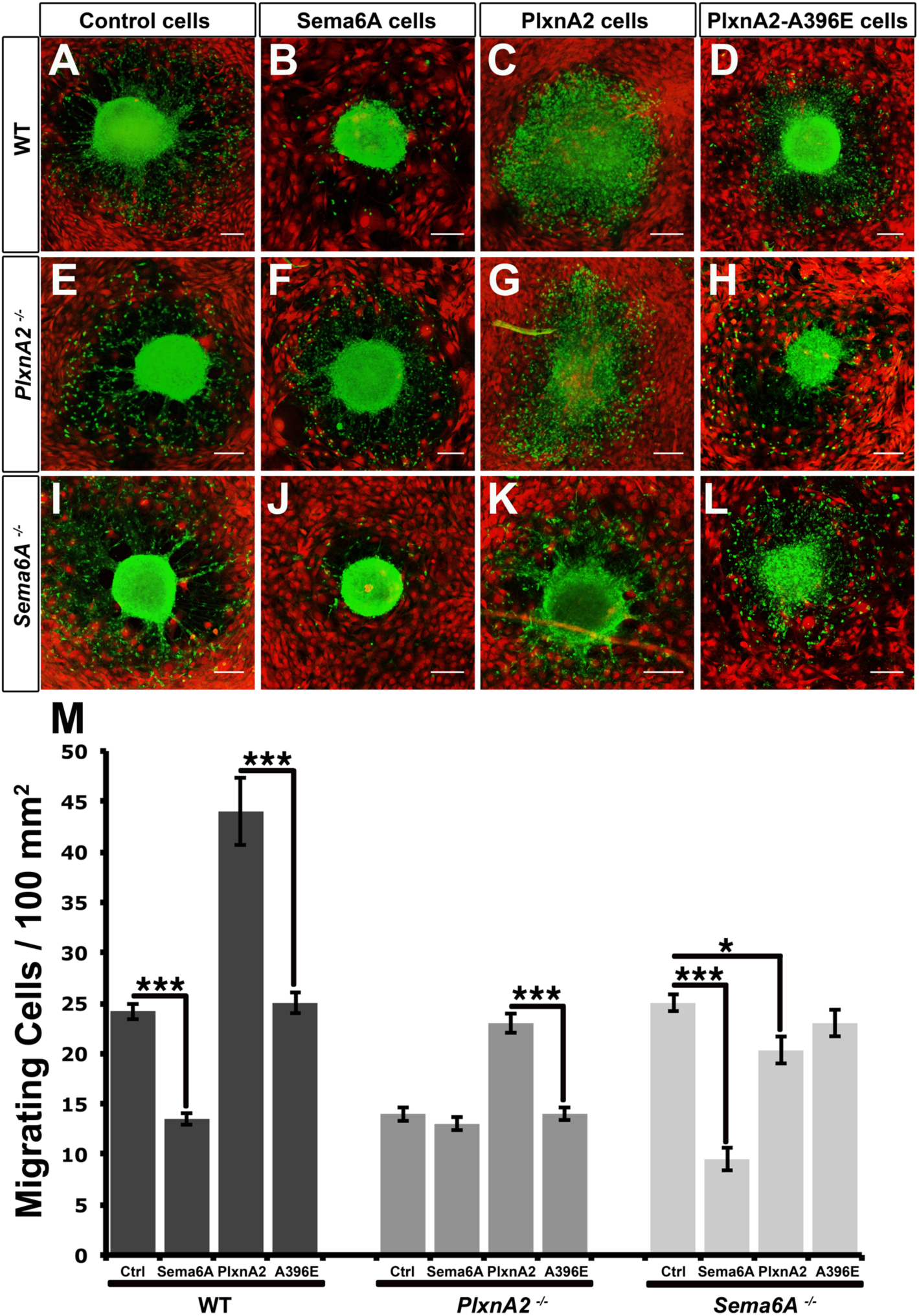
Forward and reverse Sema6A-PlxnA2 interactions have different effects on the migration of cerebellar neurons. **(a-l)** EGL explants of WT **(a-d)**, *PlxnA2^-/-^* **(e-h)**, and *Sema6A^-/-^* **(i-l)** mice cultured on a monolayer of NIH3T3 cells expressing dTomato alone (Control cells), or together with Sema6A-FL (Sema6A Cells), PlxnA2-FL (PlxnA2 Cells) and PlxnA2-A396E (PlxnA2-A396E Cells). Migrating granular neurons are visualised in green and NIH3T3 cells in red. Scale bar = 100 μm. (**m**) Graph summarising the migration of different EGL explants cultured together with different NIH3T3 layers; *n=* 50–100 explants per experimental condition. Data are expressed as mean ± s.e.m ; \**P* < 0.05, \*\**P* ≤ 0.005 and \*\*\**P* < 0.001; \*\*\**P* < 0.001; one-way ANOVA followed by Bonferroni multiple comparison test.

### PlxnA2-induced Sema6A reverse signaling reduces neurite outgrowth and complexity

To explore neuronal outgrowth linked to PlxnA2-Sema6A signaling we dissociated the EGL layer and seeded neurons onto sheets of NIH3T3 cells (Figure 3a-d). Granule neurons cultured on top of either Sema6A- or PlxnA2-expressing cells exhibited a dramatic reduction in length of the axon and branches per axon, compared to neurons cultured on untransfected or PlxnA2-A396E-expressing cells (Figure 3a-e; Figure S2). Responsiveness to Sema6A required PlxnA2 in the neurons (Figure 3f). In the opposite direction, *Sema6A^-/-^* neurons similarly did not show any response to PlxnA2-expressing sheets (Figure 3g). Responsiveness could be restored by transfection with Sema6A-FL (Figure 3g) but not with a truncated construct lacking the cytoplasmic domain (Sema6A-Δcyt) (Figure 3k). In fact, transfection of wild-type neurons with the Sema6A-Δcyt construct blocked response to PlxnA2, suggesting that this truncated form of Sema6A acts as a dominant-negative for Sema6A reverse signaling (Figure 3h-k).

**Figure 3.**
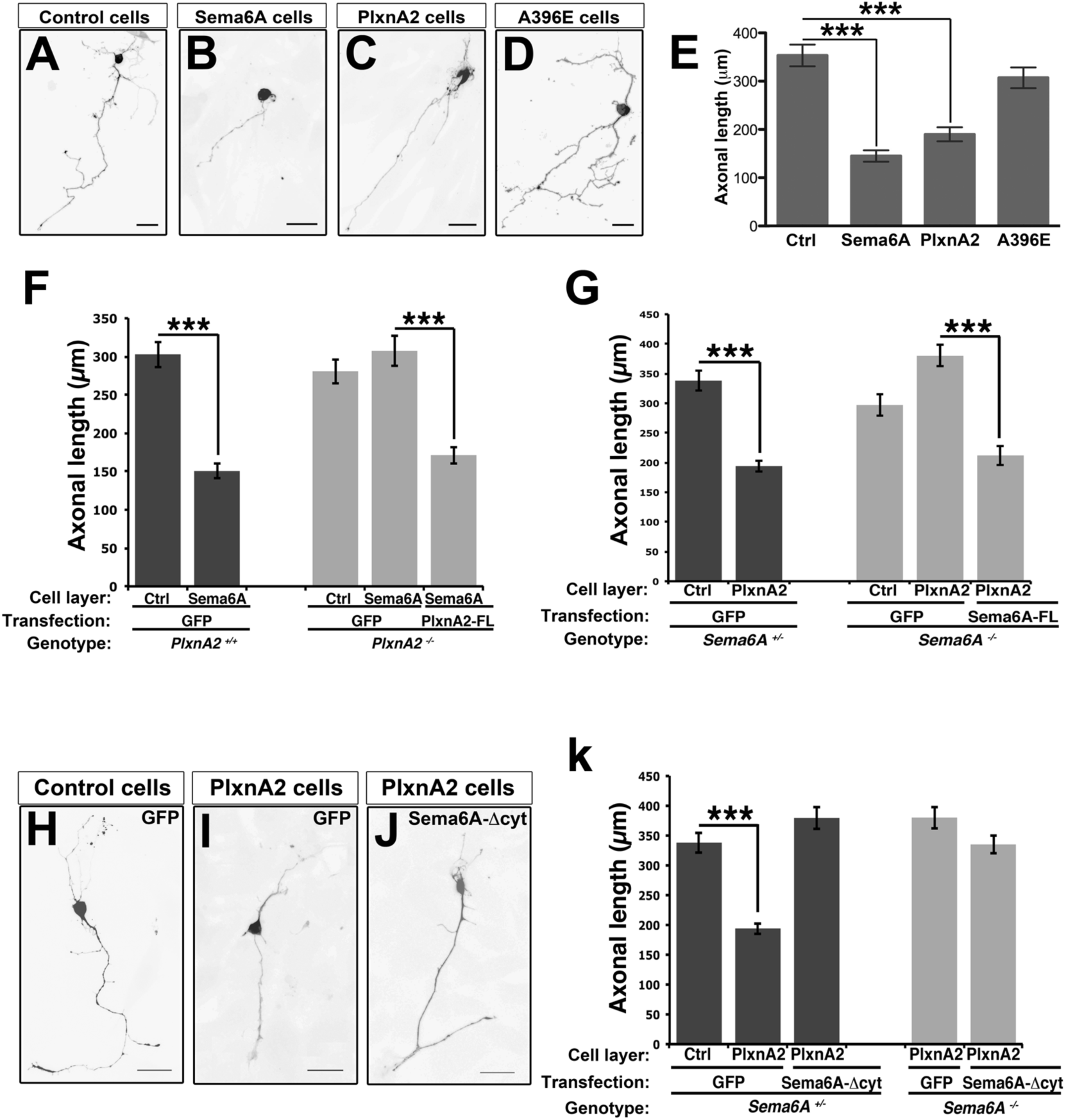
Sema6A signaling reduces the axonal length of granular neurons. (**a-c**) GFP-expressing granular neurons were cultured on Control cells (a), Sema6A Cells (**b**), PlxnA2 Cells (**c**) and PlxnA2-A396E Cells (A396E Cells, **d**). Scale bar = 20 μm. (**e**) Graph represents the axonal length of granular neurons grown on different NIH3T3 layers; *n*=100–200 cells per experimental condition. Data are expressed as mean ± s.e.m; *\*\*\*P <* 0.001; one-way ANOVA followed by Bonferroni multiple comparison test. (**f**) Graph represents the axonal length of *PlxnA2^+/+^*(dark columns) and *PlxnA2^-/-^* (light columns) granular neurons grown on NIH3T3 cells. Neurons were transfected with GFP alone or together with PlxnA2-FL, and cultured on control (Ctrl) or on Sema6A cell layers; n=100–200 cells per experimental condition. Data are expressed as mean ± s.e.m; \*\*\**P* < 0.001; Student's *t*-test. (**g**) Graph represents the axonal length of *Sema6A^+/-^* (dark columns) and *Sema6A^-/-^* (light columns) granular neurons grown on NIH3T3 cells. Neurons were transfected with GFP alone or together with Sema6A-FL, and cultured on control (Ctrl) or on PlxnA2 cell layers; *n*=100–200 cells per experimental condition. Data are expressed as mean ± s.e.m; \*\*\**P* < 0.001; Student’s *t*-test. (**h-j**) Granular neurons transfected with GFP alone (**h and i**) or together with Sema6A-Δcyt (**j**), and grown on control or on PlxnA2 cells. Scale bar = 20 μm. (k) Graph represents the axonal length of *Sema6A^+/^'* (dark columns) and *Sema6A^/-^* (light columns) granular neurons. Neurons were transfected with GFP alone or together with Sema6A-Δcyt, and cultured on control (Ctrl) or on PlxnA2 cell layers; *n*=100–200 cells per experimental condition. Data are expressed as mean ± s.e.m; ***P < 0.001; Student’s *t*-test.

### PlxnA2 can bind Sema6A in *trans* and in *cis*

Using transfected COS-7 cells, we reproduced previous findings [3] that the extracellular domain of Sema6A fused to alkaline phosphatase (Sema6A-AP) can bind to PlxnA2-FL-expressing cells but not to cells expressing PlxnA2-A396E (Figure 4a-d and f). Intriguingly, this binding was dramatically reduced when Sema6A was also co-expressed in the COS-7 cells (Figure 4e), similar to results reported for PlxnA4 [13]. By contrast, binding of PlxnA2-AP to cells expressing Sema6A-FL was not affected by co-expression of PlxnA2-FL in the COS-7 cells (Figure 4g-k). This difference is not due to a change in expression levels of the individual proteins as these are not affected by co-expression of the other protein (Figure S3).

**Figure 4.**
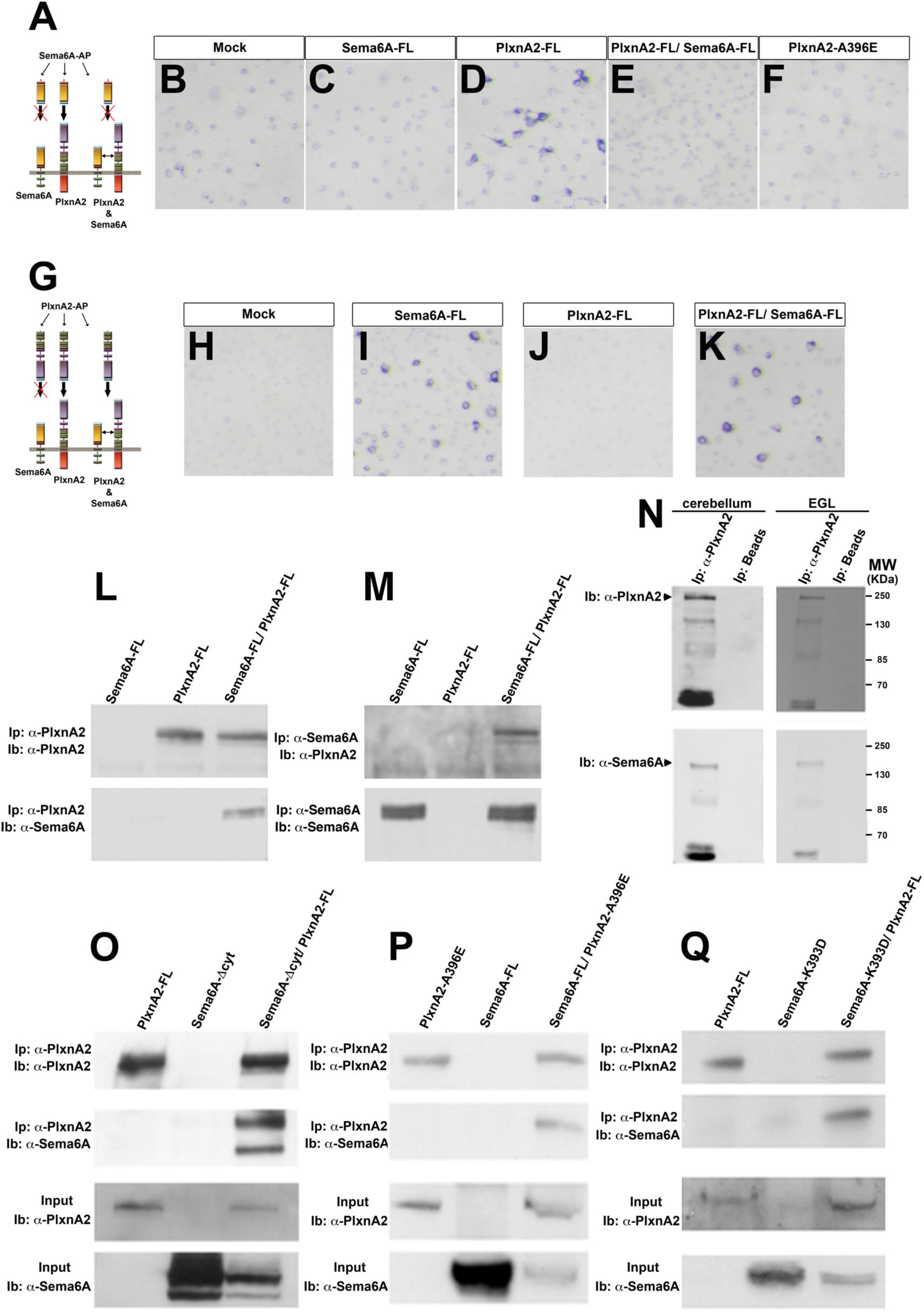
PlxnA2 interacts with Sema6A *in cis* and *in trans*. (**a**) Scheme summarising Sema6A-AP bindings. (**b-f**): COS-7 cells were transfected with the empty vector (mock; **b**), Sema6A-FL (**c**), PlxnA2-FL (**d**), PlxnA2-FL together with Sema6A-FL (**e**) or PlxnA2-A396E (**f**); and treated with Sema6A-AP. (**g**) Scheme summarising PlxnA2-AP bindings. (**h-k**): COS-7 cells were transfected with the empty vector (mock; **h**), Sema6A-FL (**i**), PlxnA2-FL (**j**) or PlxnA2-FL together with Sema6A-FL (**k**) and treated with PlxnA2-AP. (**l**) PlxnA2 immunoprecipitation (IP) from protein samples of COS-7 cells transfected with Sema6A-FL, PlxnA2-FL or Sema6A-FL together PlxnA2-FL. Antibodies against PlxnA2 and Sema6A were used in the immunoblots. (**m**) Sema6A IP from protein samples of COS-7 cells transfected with Sema6A-FL, PlxnA2-FL or Sema6A-FL together with PlxnA2-FL. (**n**) PlxnA2 IP from protein samples of mouse cerebellums or EGL explants. (**o**) PlxnA2 IP from protein samples of COS-7 cells transfected with PlxnA2-FL, Sema6A-Δcyt or Sema6A-Δcyt together with PlxnA2-FL. (**p**) PlxnA2 IP from protein samples of COS-7 cells transfected with PlxnA2-A396D, Sema6A-FL or Sema6A-FL together with PlxnA2-A396D. (**q**) PlxnA2 IP from protein samples of COS-7 cells transfected with PlxnA2-FL, Sema6A-K393D or Sema6A-K393D together with PlxnA2-FL.

To explore a possible *cis* interaction between Sema6A and PlxnA2 we performed co-immunoprecipitations from homogenates of COS-7 cells co-transfected with Sema6A-FL and PlxnA2-FL. Full-length Sema6A and PlxnA2 could be co-precipitated in pull-down experiments employing either anti-Sema6A or anti-PlxnA2 antibodies (Figure 4l and m). This interaction was also demonstrated between the endogenous proteins *in vivo* as Sema6A could be co-immunoprecipitated with PlxnA2 from homogenates of cerebellar tissue and EGL explants (Figure 4n). Considering that the Sema6A-PlxA4 *cis-* interaction takes place through the SEMA domains [13] we also performed pull-downs using cells co-transfected with PlxnA2-FL and Sema6A-Δcyt, which confirmed that the ectodomains of Sema6A and PlxnA2 were sufficient for the interaction in *cis* (Figure 4o). To elucidate whether the amino acid residues involved in the cis-interaction overlapped with the ones for the *trans-interaction* we also carried out pull-down experiments from cells co-transfected with either Sema6A-FL and PlxnA2-A396E or Sema6A-K393D and PlxnA2-FL. In both cases, *cis* interactions were still observed (Figure 4p and q). Pull-downs from cells combining the expression of PlxnA2-FL with the mutant Sema6A-I322E, which interrupts the formation of the Sema6A homodimer [29], also demonstrated that Sema6A monomer was sufficient to interact in *cis* with PlxnA2 (Figure S4).

### Sema6A cytoplasmic domain binds Abl and Mena

The Sema6A cytoplasmic domain has previously been shown to bind to Ena/VASP-like protein, Evl, through a C-terminal motif [26]. Both Sema6D and Sema1a have been shown to bind to or signal through Enabled or mouse Enabled (Mena) [20,22]. Sema6A also has a strongly predicted Abl-binding motif [27], which is in a conserved position with the demonstrated Abl-binding motif of Sema6D [22] (Figure S5a). To evaluate the binding of Abl and Mena with Sema6A we carried out a series of pull-downs from co-transfected COS7 cells, with or without stimulation with extracellular domain of PlxnA2. Abl was co-precipitated with Sema6A-FL in co-transfected cells, but only when treated with PlxnA2-EC (Figure 5a). Abl protein was also co-immunoprecipitated from homogenates of cerebellar tissue using an antibody against endogenous Sema6A, confirming this interaction in vivo and consistent with the exposure in vivo to co-expressed PlxnA2 in cerebellar granule cells (Figure S5b). By contrast, Mena was pulled down together with Sema6A in cells either untreated or treated with PlxnA2-EC (Figure 5b) and also from cerebellar extracts (Figure S5c).

**Figure 5.**
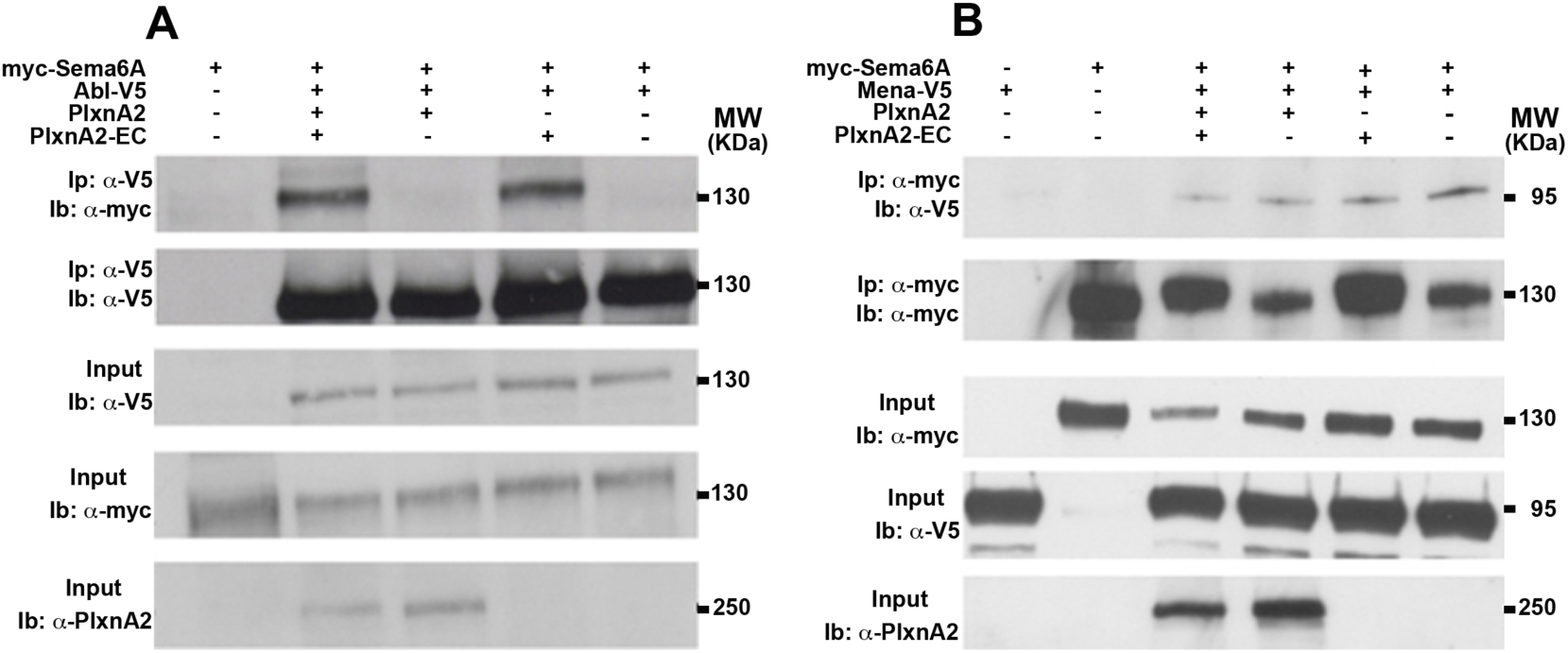
Sema6A cytoplasmic domain interacts with Abl and Mena. (**a**) Immunoprecipitations from untreated or PlxnA2-EC-treated COS-7 cells previously transfected with different combinations of myc-Sema6A, Abl-V5 and PlxnA2. Immunoprecipitations (Ip) were performed employing a α-V5 antibody. Antibodies against V5, myc and PlxnA2 were used in the immunoblots (Ib). (**b**) Immunoprecipitations from untreated or PlxnA2-EC treated COS-7 cells previously transfected with different combinations of myc-Sema6A, Mena-V5 and PlxnA2. Immunoprecipitations (Ip) were performed employing a α-myc antibody. Antibodies against V5, myc and PlxnA2 were used in the immunoblots (Ib).

### Sema6A reverse signaling is stimulated by multimerisation

The crystal structure of the Sema6A-PlxnA2 interaction suggested that formation of higher-order clusters might be important for activating signaling [29,30]. To explore this possibility, we designed a chimeric construct putting in frame three self-aggregation FKBP domains [34], the entire cytosolic tail of Sema6A (Sema6A-cyt) and EGFP as a reporter. We also included a myristoylating sequence at the N-terminus of the chimeric peptide to direct anchorage of Sema6A to the plasma membrane (Figure 6a). Using this system we were able to induce the clustering of Sema6A-cyt in the presence of FK1012, an inert and membrane-permeable chemical, which has bivalent interactions with the FKBP domain [34] (Figure 6a). Cerebellar granular neurons were transfected with either a control construct (MF3-EGFP) or the Sema6A-cyt construct (MF3-Sema6A-cyt). The EGFP reporter initially showed diffuse staining throughout the cell body and neurites of transfected neurons. This signal became dramatically clustered in response to incubation with 500nM FK1012 for 30 min (Figure 6b). We compared the FK1012-inducible multi-aggregation of Sema6A-cyt with the incubation of PlxnA2-EC on top of neurons transfected with myc-Sema6A-FL. We observed an increase in myc-Sema6A-FL clustering in neurons treated with PlxnA2-EC comparable to the FK1012-inducible Sema6A-cyt-EGFP (Figure 6c and d). The *trans-interaction* with PlxnA2-EC thus promotes the *in situ* clustering of Sema6A in the plasma membrane (Figure 6d). Finally, we compared the axonal length of untreated vs FK1012-treated neurons previously nucleofected with MF3-Sema6Acyt and cultured on top of untransfected NIH3T3 sheets. The resultant measurements revealed that induction of Sema6A-cyt clustering by chronic chemically induced multimerisation was enough to elicit a significant reduction of the axonal length, similar to the effects of exposure to PlxnA2 (Figure 6e and f).

**Figure 6.**
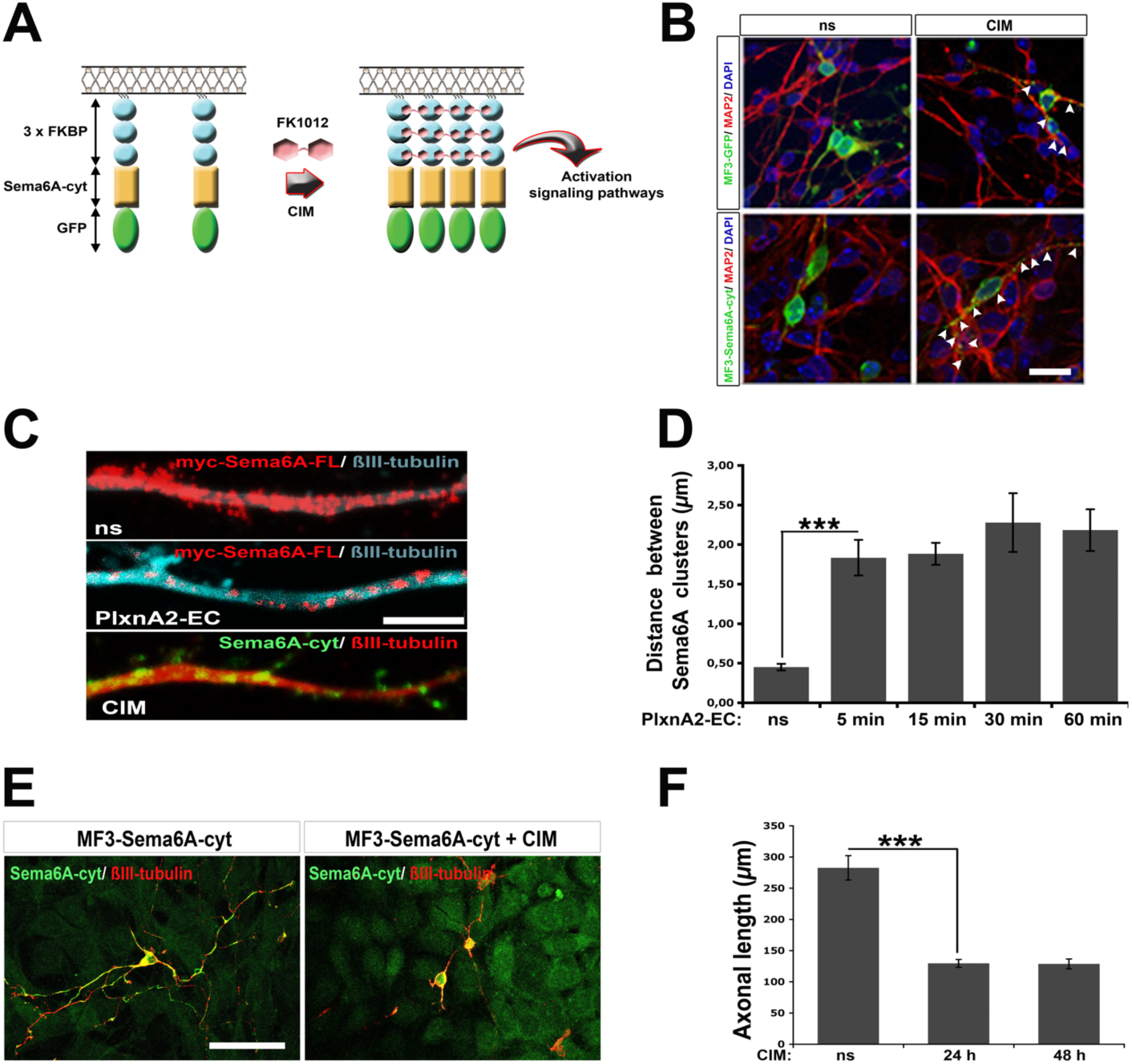
Sema6A reverse signaling is stimulated by multimerisation. (**a**) Scheme representing how the induced multi-aggregation of the cytosolic domain of Sema6A activates signaling pathways. A chimeric construct was engineered by putting in frame the entire cytosolic domain of Sema6A (Sema6A-cyt) together with 3 self-aggregation FKBP domains, and GFP. The chimeric peptide targets the plasma membrane by the myristoylating sequence located at the N-terminal. The application of the drug FK1012 triggers the chemically induced multimerisation (CIM) of Sema6A-cyt via the self-aggregation of FKBP. (**b**) Cerebellar neurons transfected with MF3-GFP or MF3-Sema6A-cyt-GFP (MF3-Sema6A-cyt), and treated with 500nM FK1012 (“CIM”). GFP-positive aggregates are indicated (arrowheads). Scale bar = 20μm. (**c**) Axonal shaft of cerebellar neurons transfected with myc-Sema6A-FL or MF3-Sema6A-cyt-GFP (MF3-Sema6A-cyt), and treated with PlxnA2-EC or CIM respectively. Scale bar = 5 μm. (**d**) Graph summarising the myc-Sema6A-FL aggregation in myc-Sema6A-FL-expressing neurons treated with PlxnA2-EC at different time points. The aggregation is represented as the average distance between Sema6A clusters; *n*=50–100 cells per time point. Data are expressed as mean ± s.e.m; \*\*\**P* < 0.001; one-way ANOVA followed by Bonferroni multiple comparison test. (**e**) MF3-Sema6A-cyt-expressing granular neurons were cultured on NIH3T3 cell layers, and treated with CIM. Scale bar = 50 μm. (**f**) Graph represented the axonal length of MF3-Sema6A-cyt-expressing cerebellar neurons treated with CIM for 24 and 48h; *n*=50–100 cells per experimental condition. Data are expressed as mean ± s.e.m; \*\*\**P* < 0.001; one-way ANOVA followed by Bonferroni multiple comparison test.

### Sema6A multimerisation stimulates phosphorylation of cytoskeletal regulators

To evaluate the potential signaling pathways initiated by Sema6A clustering, we screened a panel of proteins for phosphorylation after the chemically induced multimerisation of Sema6A-cyt in MF3-Sema6A-cyt-transfected neurons. These proteins were chosen to represent major signal transduction pathways or based on known interactions with the cytoskeleton.

Acute induction of chemically induced multimerisation (from 15 to 45 min) increased the phosphorylation of the kinases Abl and GSK3α/β as well as the actin-adaptor proteins p130CAS, Ezrin/Radixin/Moesin and MARCKS (Figure S6a). These results were consistent across three independent replicates. We also observed increased phosphorylation of p130Cas by immunocytochemistry and a tight spatial overlap of phospho-p130CAS signal with Sema6A-cyt clusters (Figure S6b), providing additional evidence of the linkage between the *in situ* multimerisation of Sema6A-cyt with actin-related signaling cues. By contrast, multimerisation of Sema6A-cyt did not stimulate the MAP kinase pathway or lead to PAK1 or CREB phosphorylation (Figure S6a).

In accordance with the signaling results obtained from chemically induced multimerisation of Sema6A-cyt we evaluated whether the stimulation of neurons with PlxA2-EC could also trigger actin-signaling cues, using p130CAS and MARCKS as test examples. Granular neurons exhibited an increase of the phosphorylation of both these proteins after 15 minutes of incubation with PlxnA2-EC at a level comparable to the chemically induced multimerisation Sema6A-cyt activation (Figure S6c).

### PlxnA2 responsiveness in NIH3T3 cells requires the Abl-binding domain

To assess the functional relevance of the Abl signaling pathway, we designed an expression construct of full-length Sema6A with the putative Abl-binding motif deleted, Sema6A-ΔAbl (Figure S5d). This construct is unable to bind to Abl in co-immunoprecipitation assays (Figure S5e). Transfection of this construct into NIH3T3 cells produced the same expansion of cellular area and filopodial extensions observed with wild-type Sema6A (Figure 7a-c). However, it did not confer responsiveness to PlxnA2 (Figure 7d-h).

**Figure 7.**
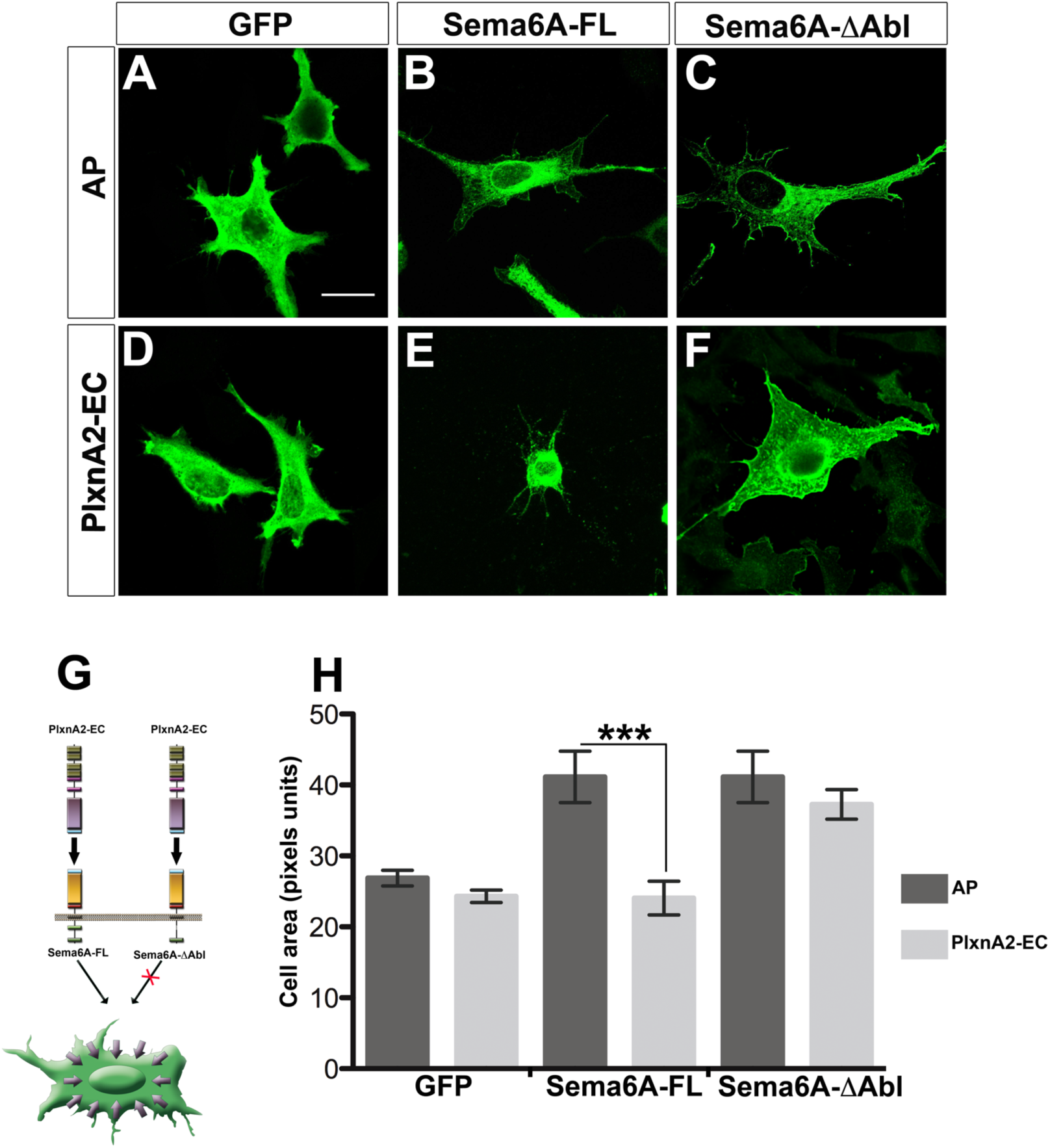
Cell-contraction induced by PlxnA2-Sema6A interaction depends on Abl signaling. **(a-f)** NIH3T3 cells expressing GFP **(a and d)**, Sema6A-FL **(b and e)**, and Sema6A-AAbl **(c and f)** were treated with purified AP-Fc (AP; **a-c)** or PlxnA2-EC-Fc (PlxnA2-EC; **d-f)**. Scale bar = 20 μm. **(g)** Scheme illustrates that the cell contraction is via Abl signaling, **(h)** Graph represents the contraction of the cell area in NIH3T3 transfected with GFP, Sema6A or Sema6A-Mbl and treated with AP or PlxnA2-EC; *n*=100–200 cells per experimental condition. Data are expressed as mean ± s.e.m; \*\*\**P* < 0.001; Student's *t*-test.

## Discussion

Our findings demonstrate that Sema6A can engage in reverse (cell-autonomous) signaling, constitutively and in response to either ligand-binding or artificially induced multimerisation. This signaling induces recruitment and phosphorylation of Abl tyrosine kinase and the phosphorylation of additional cytoplasmic signaling proteins, many involved in cytoskeletal regulation. Activation of Sema6A has multiple effects on non-neuronal and neuronal cells *in vitro*, influencing cell morphology, neurite growth and cellular aggregation.

In NIH3T3 fibroblast cells, expression of full-length Sema6A induces a change in cell shape, with an increase in surface area and in membrane complexity, with numerous filopodia apparent. It also confers responsiveness to the extracellular domain of PlxnA2. PlxnA2 has no effect on untransfected or GFP-transfected cells but induces cellular collapse of cells transfected with full-length Sema6A. By contrast, cells transfected with a mutant form of Sema6A that does not bind PlxnA2 show the increase in cellular area and complexity but no response to PlxnA2. These data suggest that not only can Sema6A initiate reverse signaling – it can do it in two distinct modes. The fact that the effects of stimulated and unstimulated signaling are qualitatively distinct, perhaps even opposite to each other, argues against the idea that constitutive signaling is merely due to over-expression. We speculate that these two modes may reflect signaling by unclustered dimers of Sema6A versus multimerised clusters induced by ligand binding.

We used an assay with cerebellar granule cell explants to assess Sema6A signaling in a more physiological context. Our data clearly demonstrate that activation of endogenous Sema6A in neurons, by exposure to PlxnA2, stimulates two distinct effects. First, explants grown on NIH3T3 cells expressing PlxnA2 are strongly disaggregated, an effect that depends on Sema6A activity in the responding cells. This disaggregation likely reflects an alteration of paracrine interactions between cerebellar granule cells. If Sema6A activation mediated a typical repulsive signal, one might expect a tighter aggregation of the explant as Sema6A-positive cells avoided the PlxnA2-expressing substrate. Instead, exposure to PlxnA2 from the substrate alters interactions *between* cerebellar granule cells, suggesting that Sema6A activation may feed back to the cell surface, regulating mutual repulsion or levels of adhesion.

Second, in dissociated cerebellar granule neurons, activation of Sema6A signaling has effects on neurite outgrowth and complexity. Wild-type neurons grown on sheets of PlxnA2-expressing cells have reduced axonal and dendritic length and number of branches. *Sema6A^-/-^* neurons are insensitive to PlxnA2, but transfection of a construct encoding full-length Sema6A restores responsiveness. The fact that a construct of Sema6A lacking the cytoplasmic domain not only cannot signal but acts as a dominant-negative further reinforces the conclusion that these effects involve signaling through the Sema6A cytoplasmic domain. This is directly demonstrated by the fact that chemically induced multimerisation of the membrane-tethered Sema6A cytoplasmic domain is sufficient to induce similar effects on neurite outgrowth in dissociated neurons.

Our biochemical assays have identified a number of downstream events following Sema6A activation, involving phosphorylation of multiple proteins with functions in cytoskeletal regulation. The most direct interaction is with the cytoplasmic tyrosine kinase, Abl, which acts as an intermediary between transmembrane receptors and the cytoskeleton, affecting cellular morphology, adhesion and migration [35]. Sema6A activation by ligand binding leads to recruitment of Abl kinase to the cytoplasmic domain and to Abl phosphorylation. We show, using a deletion construct, that Abl recruitment is required for PlxnA2-induced cellular collapse of NIH3T3 cells, suggesting it is a proximal event in Sema6A ligand-induced reverse signaling, though this role remains to be demonstrated directly in neurons. It is not however required for the increase in cellular area and complexity that accompanies full-length Sema6A expression, providing additional evidence for a biochemically distinct, constitutive mode of reverse signaling by Sema6A.

The Sema6A cytoplasmic domain also directly interacts with the Abl substrate Mena, in a manner that is independent of ligand binding. The functional relevance of this interaction remains to be tested, but an important role in regulating adhesion or migration would be consistent with known functions of Ena/VASP proteins [36] and with the implication of both Abl and Enabled orthologues in reverse signaling by Sema6D and Sema1a [20,22]. The Rho signaling modulators p190RhoGAP and pebble (a RhoGEF) have also recently been implicated in Sema1a reverse signaling in Drosophila, through direct interactions with the cytoplasmic domain [21]. Interestingly, p190RhoGAP is a substrate for the Abl orthologue, Arg, suggesting a possible interaction between these signaling pathways to regulate Rho1 activity and coordinate effects on the cytoskeleton or on adhesion [37,38,39].

Sema6A activation also triggers phosphorylation of a number of other cytoskeleton-regulating cytoplasmic proteins, including p130CAS [40], MARCKS [41], ezrin, radixin and moesin [42] and GSK3-alpha [43,44]. These molecules have well-documented effects on cellular morphology, adhesion, neurite outgrowth, branching or guidance (see reviews referenced above). Many of these pathways have known interactions with Abl signaling in regulation of actin-cytoskeletal rearrangements with diverse cellular effects [45,46,47,48,49,50]. However, the functional roles and relationships of these various proteins in the cellular events induced by Sema6A reverse signaling remain to be determined.

In addition to identifying functional bidirectional signaling arising from Sema6A-PlxnA2 *trans* interaction, we also observe interactions *in cis* between these molecules (Figure 8). These are mediated by the extracellular domains but do not depend on the same interfaces as *trans* interactions, as amino acid mutations that block *trans* interactions leave *cis* interactions intact. Nevertheless, the *cis* interactions can interfere with the *trans* interactions: co-expression of Sema6A *in cis* blocks the ability of PlxnA2 to bind Sema6A-EC in *trans*. Evidence for a similar effect was recently observed *in vivo* in the developing retina [12], and it is similar to the previously described effect on PlxnA4 [13]. The converse was not observed however; Sema6A could bind PlxnA2-EC in *trans* and Abl could be consequently recruited to the Sema6A cytoplasmic domain, regardless of the presence or absence of PlxnA2 in *cis*. In the opposite direction, co-expression of PlxnA2 in *cis* has been proposed to disrupt Sema6A function as a ligand, with PlxnA2 expression masking the repulsive effects of Sema6A and Sema6B and restricting hippocampal mossy fiber projections to a specific layer [10,11]. A similar role for PlxnA2 in masking a repulsive Sema6A signal has been reported in the developing eye vesicle in zebrafish [51].

**Figure 8.**
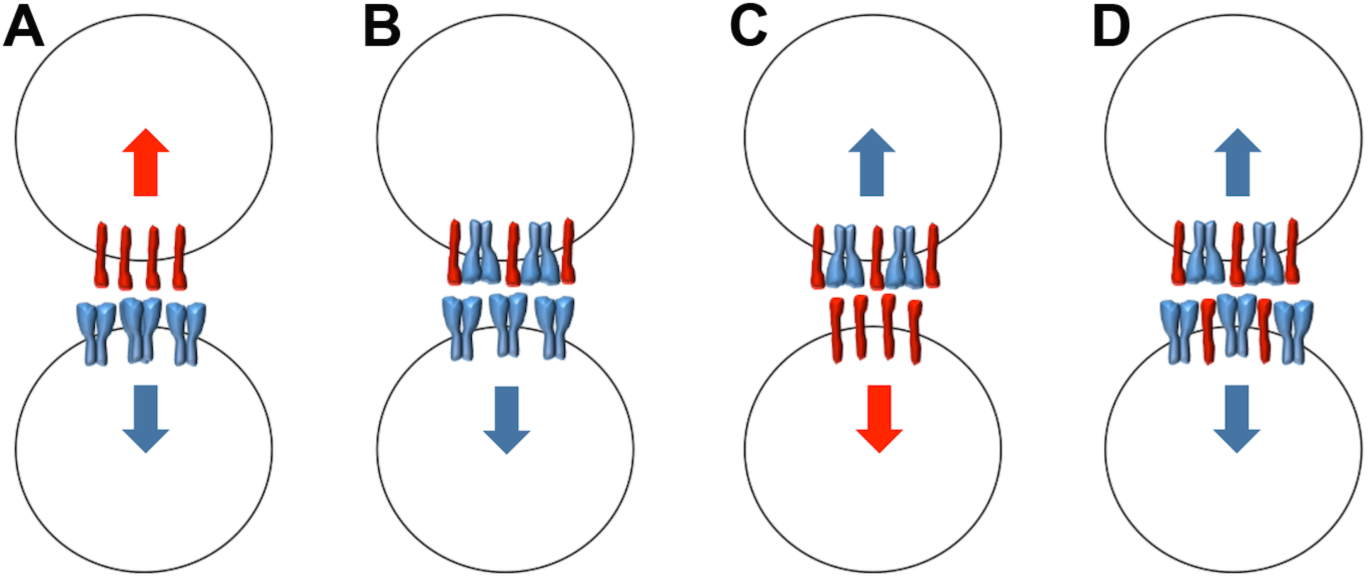
Schematic of diverse Sema6A-PlxnA2 interactions. Neighbouring cells are shown expressing either Sema6A (blue), PlxnA2 (red) or both molecules. (a) Signaling between cells expressing Sema6A alone and PlxnA2 alone can proceed in both directions (blue and red arrows). (b) Expression of Sema6A *in cis* blocks receptor function of PlxnA2 in the top cell but PlxnA2 can still act as a ligand to stimulate Sema6A receptor function in the bottom cell (blue arrow). (c) Sema6A-PlxnA2 complexes can act as receptors for PlxnA2 *in trans* (top cell), but not for Sema6A *in trans* (top cell in b). In some cases *in vivo* [12], Sema6A can still act as a ligand when co-expressed with PlxnA2 (red arrow, bottom cell). In others [10,11,51], PlxnA2 may directly occlude or indirectly antagonise this function (scenario not shown). (d) In populations of cells expressing both Sema6A and PlxnA2, the complexes should act as receptors for PlxnA2, through reverse signaling by Sema6A (blue arrows). The balance between these various interactions will likely depend on the precise stoichiometry of the proteins and may also be dynamically altered by interactions with additional proteins or newly encountered cells (e.g., ref. 53).

Taken together, these findings suggest that heteromeric Sema6A-PlxnA2 complexes should present PlxnA2 (but not Sema6A) as a ligand to other cells but should function as receptors through reverse signaling by Sema6A (but not through PlxnA2) (Figure 8). A recent study of the functions of these genes in the retina supports this model [12]. Sema6A and PlxnA2 are co-expressed in ON starburst amacrine cells (SACs) and both are required for dendritic self-avoidance. The authors confirm that Sema6A is repulsive for neurites of SACs expressing PlxnA2 alone (mediating segregation of ON and OFF SAC dendrites) but not SACs expressing both PlxnA2 and Sema6A. If PlxnA2 cannot act as a receptor when Sema6A is expressed *in cis*, this suggests instead that Sema6A may function as the signaling receptor for self-avoidance, responding to PlxnA2 as the ligand in neighboring dendrites.

More direct evidence of Sema6A reverse signaling in vivo was very recently described by the same authors, who showed that Sema6A is required in a subset of retinal ganglion cells as a receptor for PlxnA2 and PlxnA4, to promote correct innervation by these axons of the medial temporal nucleus (MTN) [52].

Finally, a recent study in the developing spinal cord illustrates a similar scenario, in this case involving Sema6B and PlxnA2 [53]. Commissural neurons express Sema6B, which acts as a receptor for PlxnA2 expressed by the floorplate, with this interaction promoting midline crossing. PlxnA2 is co-expressed in cis by commissural neurons but does not block the response to exogenous PlxnA2. However, PlxnA2 is prevented by the cis interaction with Sema6B from responding prematurely to the secreted semaphorin Sema3A. When Sema6B binds PlxnA2 from the floorplate in trans, this cis interaction is disrupted and PlxnA2 becomes an active receptor for Sema3A, which ensures accurate post-crossing projections.

The findings we present here provide a biochemical framework for the mechanisms underlying these complex and context-dependent interactions. The Sema6 and PlxnA proteins do not fall into neat ligand-receptor categories but exhibit a much more intricate and combinatorial molecular logic, often simultaneously mediating multiple cellular interactions: between different cell types, among groups of cells or nerve fibres through paracrine interactions or even across neurites from the same cell. The biochemical activities we have defined will inform the interpretation of mutant phenotypes for these genes in the expanding list of regions where they play important roles [1].

## Materials and Methods

All animal work was approved by the Ethics Committee of Trinity College Dublin and the animal licence to KJM (Ref. B100/3527) from the Department of Health and Children.

### Constructs

PCAGGS-myc-Sema6A-FL, pCAGSS-flag-Sema6A-FL, pCAGSS-flag-Sema6B-FL, pCAGGS-flag-Sema6D-FL, pCAGGS-myc-PlxnA2-FL, pCAGGS-AP-Fc-(His)6, pCAGSS-Sema6A-EC-AP-Fc-(His)6 and pCAGGS-PlxnA2-EC-AP-Fc, pCAGGS-PlxnA2-EC-Fc-(His)_6_, were kindly provided by Professor H. Fujisawa. PlexinA2-A396E was cloned into pCDNA3.1 vector as was previously described [3]. cDNAs encoding myc-tagged Sema6A-K393D, myc-tagged Sema6A-I322E, myc-tagged Sema6A-Δcyt, myc-tagged Sema6A-ΔAbl, and dTomato were cloned into pEF1 vector. Cytosolic domain of Sema6A (Sema6A-cyt) was amplified by PCR and cloned along EGFP within pC4MFv2E vector as part of the inducible homodimerisation system (Clontech). The plasmid pEGFP (Clontech) was employed to transfect neurons. Full-length V5-tagged Abelson kinase (Abl-V5) and full-length V5-tagged Mena (Mena-V5) were kindly provided by Dr. T. Toyofuku.

### Antibodies and reagents

Antibodies anti-myc (clone 9E10; mouse; 1:4000) and anti-flag (clone M2; mouse; 1:500) were supplied by Sigma-Aldrich, anti-V5 (mouse; 1:1000) by Invitrogen, and anti-AP (alkaline phosphatase; rabbit; 1:6000) antibody by GenHunter. Neuronal markers anti-NeuN (hexaribonucleotide binding protein-3; mouse; 1:200) and anti-MAP2 (microtubule associated protein-2; rabbit; 1:250) were obtained from Millipore. Anti-phospho-(S152/ S156)-MARCKS (myristoylated alanine-rich C-kinase substrate; rabbit; 1:1000), anti-phospho-(Y165)-p1030CAS (CAS scaffolding protein family member 1; rabbit; 1:200 −1:1000), anti-phospho-(S133)-CREB (cAMP response element-binding protein; rabbit; 1:1000), anti-phospho-(T567) Ezrin/ (T564) Radixin/ (T558) Moesin (rabbit; 1:1000), anti-phospho-(T202/Y204)-MAPK (mitogen-activated protein kinase; rabbit; 1:1000) were purchased from Cell Signaling; and anti-phospho-(T212)-PAK1 (protein kinase-1; rabbit; 1:500), anti-phospho-(S21/ S9)-GSK3α/β (glycogen synthase kinase-3; rabbit; 1:1000), anti-phospho-(Y245)-cAbl (Abelson leukemia protein; rabbit; 1:500) and anti-Hsp90 (heat shock protein-90; mouse; 1:1000) from Abcam. Anti-βIII-tubulin (mouse; 1:2000) was purchased from Promega. Anti-PlexinA2 (clone C18; goat; 1:250; Santa Cruz) and anti-Sema6A (goat; 1:250; R&D) were employed to detect and precipitate PlxnA2 and Sema6A. Phalloidin-rhodamin (1:500; Invitrogen) was used to stain F-actin network. FK1012 (Clontech) was used to induce artificial multimerisation.

### Animals

Wild Type (WT), PlexinA2 knock out *(PlxnA2^-/-^)* and Semaphorin6A knock out *(Sema6A^-/-^)* C57BL/6 mice at postnatal days 5–7 (P5-P7) were used. Animals were sacrificed according to Agriculture Irish Department and European Ethical and Animal Welfare regulations.

### COS-7, NIH3T3 and HEK293 cell cultures

HEK293, COS-7 and NIH3T3 were cultured in Dulbecco’s Modified Eagle Medium (DMEM) supplemented with 10% fetal bovine serum (FBS) and L-glutamine in a 5% CO_2_, 95% humidity incubator at 37°C.

### Creation of NIH3T3 stably transfected cell lines

NIH3T3 cells were transfected with dTomato alone or together with flag-Semaphorin6A-FL, myc-PlxnA2-FL or PlxnA2-A396E. After three rounds of selection, cell clones were chosen based on two criteria: 1 - The co-expression of both genes and 2 - The localization of Sema6A and PlxnA2 in the plasma membrane. Both criteria were tested by immunofluorescent techniques (IF).

### Explant and neuronal cultures

Cerebella from mice at P5 were dissected and cut into 200μm thick slices. Next, the external granular layer (EGL) was isolated from the cerebellar cortex and cut into 200 μm pieces. EGL explants were cultured on monolayers of NIH3T3 cells expressing different exogenous genes (dTomato, flag-Sema6A-FL/dTomato, myc-PlxnA2-FL/dTomato, PlxnA2-A396E/dTomato). Co-cultures were cultured in DMEMK (DMEM supplemented with L-glutamine, 10% FBS and 25mM KCl), for 4 days in 5% CO_2_, 95% humidity incubator at 37°C. Granular neurons (GN) were obtained from dissociated EGL layers, transfected, and then grown either on surfaces coated with poly-L-ornithine and fibronectin (Sigma-Aldrich) or on monolayers of transfected NIH3T3 (see above). GN and GN-NIH3T3 were cultured in DMEMK for 1–4 days in 5% CO_2_, 95% humidity incubator at 37 °C.

### Transfection in cell lines and neurons

NIH3T3 cells were transfected using polyethylene protocol (PEI). Cells were seeded the day before transfection at 1 × 10^5^ cells/ cm^2^. Thereafter, 0.5–2.5 μg of DNA were mixed up with PEI diluted in 150mM NaCl solution and incubated for 15–30 minutes (min) at room temperature (RT) to facilitate the formation of DNA-PEI complexes. Next, DNA-PEI complexes were incubated with the cells for 24–48 hours (h) in 5 % CO_2_, 95 % humidity incubator at 37 °C. HEK293 and COS-7 cells were transfected with Lipofectamin 2000 according to manufacturer instructions (Invitrogen). GN were transfected using Amaxa Nucleofection System according to manufacturer specifications (Lonza).

### Contraction Assay

60–100nM of purified AP-Fc (AP) and PlxnA2-EC-Fc (PlxnA2-EC) peptides [10] were used to treat transfected NIH3T3 cells and GN. Thereafter, cells were fixed, immunolabeled and digitally analysed.

### Induced multimerisation in neurons

GN were transfected either with pC4MFv2E-GFP (MF3-GFP) or with pC4MFv2E-Sema6A-cyt-GFP (MF3-Sema6A-cyt), and then seeded at 2.5 × 10^5^ in 24-well plates, and at 2.5-5 × 10^6^ in 6-well plates. In acute assays, 24-48h after transfection, GN cultures were treated for 15–45 min at 37 °C with 500 nM FK1012 diluted in stimulating medium (SM: DMEM, Glutamax, sodium pyruvate, 50 μM 2-mercaptoethanol and 5 % FBS). Next, cells were properly harvested for further IF and immunoblotting (IB) assays. In chronic experiments, 24h after GN transfection, GN-NIH3T3 cultures were stimulated with 500 nM FK1012 in DMEM for 24–48h. FK1012 was replaced every 24h. Co-cultures were thus fixed, immunolabeled and digitally analysed.

### *In situ* binding assay

HEK293 cells were transfected with Sema6A-EC-AP-Fc (Sema6A-AP) or PlexnA2-EC-AP-Fc (PlxnA2-AP) and grown for 48h. Supernatants were harvested and filtered to obtain conditioned media. Conditioned media were validated by alkaline phosphatase (AP) assays to confirm AP activity (Fouquet, 2007); and by Ib assays blotted with an anti-AP antibody to detect Sema6A-AP and PlxnA2-AP at the expected molecular size (data not shown). Conditioned media positive for Sema6A-AP and PlxnA2-AP were used to treat COS-7 cells previously transfected with different combinations of Sema6A and PlxnA2 constructs [3].

### Immunoprecipitation (IP)

In Sema6A and PlxnA2 pull-down assays, transfected COS-7 cells were lysed with NP-40 buffer (10mM HEPES pH 7.5; 100mM NaCl, 2mM EDTA, 0.5 % NP-40) supplemented with protease and phosphatase inhibitors, incubated at 4°C for 20 min and centrifuged at 14000 × *g* for 10 min at 4°C. Resultant supernatants were incubated for 2h at 4°C with anti-PlexinA2 and anti-Sema6A. In *in vivo* pull-down assays, lysates were prepared from P5 cerebellum or EGL explants lysed with buffer containing 50 mM Tris pH 7.6; 1mM EDTA, 1% Triton x-100 supplemented with protease and phosphatase inhibitors and incubated during 45 min rocking at 4°C. Lysates were incubated for 16h at 4°C with anti-PlexinA2.

In Abl, Mena and Sema6A-ΔAbl pull-down assays, transfected COS-7 cells were serum-starved for 16h, stimulated with freshly conditioned medium expressing PlexinA2-EC for 20 min at 37°C, and lysed. Lysates were incubated for 2h at 4°C with anti-myc or anti-V5.

All the antibody-antigen complexes were incubated with protein A/G sepharose (Amersham) for 1h at 4°C, washed with cold lysis buffer and boiled in Laemmli SDS protein sample buffer.

### Immunofluorescence (If)

EGL-NIH3T3 cultures were fixed in 4% paraformaldehyde supplemented with 0.01% Triton X-100 (PFAT) for 1h at RT. After several rinses with phosphate buffered saline (PBS) supplemented with 0.01% Triton X-100, cultures were pre-incubated with immunofluorescence buffer 1 (IFB1: PBS + 0.01% Triton X-100 and 5 % normal donkey serum) for 1h at RT, and subsequently incubated overnight (ON) at 4°C with anti-NeuN in IFB1.

GN-NIH3T3 cultures were fixed in PFAT for 15 min at RT. After several rinses with PBS supplemented with 0.01% Triton X-100, cultures were pre-incubated ON at 4°C with IFB1 for 1h at RT, and subsequently incubated with anti-βIII-tubulin in IFB1.

Transfected NIH3T3 and GN cultures were fixed in 4% paraformaldehyde (PFA) for 15 min. After several rinses with PBS, cultures were pre-incubated with immunofluorescence buffer 2 (IFB2: PBS + 0.1% Triton X-100 and 5% normal donkey serum) for 1h at RT, and subsequently incubated ON at 4°C with proper primary antibodies diluted in IFB1.

Suitable secondary antibodies conjugated to Alexa fluorochromes were used for further microscopic analysis. Moreover, F-actin was detected using phalloidin-rhodamine.

### Protein sample preparation

1 × 10^5^ HEK293, 1 × 10^5^ COS-7 and 2.5-5 × 10^6^ GNs were harvested in cold homogenization buffer (50mM Tris pH 8.0; 5mM EDTA, 140mM NaCl) supplemented with 1mM dithiothreitol (DTT) and protease and phosphatase inhibitors. Homogenates were centrifuged at 1000 × *g* for 10 min to remove nuclear fraction; and resultant supernatants (S1) were incubated with a final concentration of 1% of Triton X-100 for 30 min rocking at 4°C, and centrifuged at 11000 × *g* for 30 min at 4°C. Supernatants (S2) were thus submitted to protein quantification.

### Immunoblotting (Ib)

Protein samples from HEK293, COS-7, GN, and IPs were submitted to SDS-PAGE and blotted onto nitrocellulose membranes. Blots were incubated ON at 4°C with proper primary antibodies diluted in Tris-buffered saline and 0.1% Tween-20 (TBST) supplemented with 5% non-fatty dry milk. After several washes with TBST, blots were incubated with corresponding secondary antibodies conjugated to horseradish peroxidase (HRP), and developed by chemical luminescence techniques.

### Analysis and quantification

*In situ* binding assays on COS-7 cells were reported employing a Leica MZ16 stereoscope. Transfected NIH3T3, EGL-NIH3T3, GC-NIH3T3, transfected GN cultures were examined employing a Zeiss LSM-700 microscope (Zeiss). In EGL-NIH3T3 cultures, the migration of postmitotic cell bodies co-cultures was counted and the explant area measured by employing ImageJ. Migration data were normalized as number of migrating cells for 100 μm^2^ of explant area. 50–100 explants per each experimental condition were analysed.

Neurite length and complexity in GN-NIH3T3 cultures, the aggregation of myc-Sema6A in transfected GN, and the cell area in transfected NIH3T3 cells were analysed using ImageJ (http://rsbweb.nih.gov/ij/). 50–200 cells per experimental condition were evaluated.

Statistical analysis was performed using Prism software as indicated (GraphPad software). Data were shown as mean ± standard error of the mean (s.e.m). Statistical analysis was carried out employing the Student’s *t*-test for unpaired variables (two-tailed) and one-way ANOVA followed by Bonferroni multiple comparison tests when three or more groups were compared. *P*-values < 0.05 were considered significant.

## Acknowledgements

We thank Professor Juan Pablo Labrador, Professor Beate Winner and Professor Juergen Winkler for excellent support and critical input. We are very grateful to Dr. Jackie Dolan and members of the Trinity College BioResources Unit for their help with animal maintenance, breeding and genotyping. This work was supported by grants from Science Foundation Ireland (07/IN.1/B969 and 09/IN.1/B2614) to KJM and by grants to Alain Chédotal from the Fondation pour la Recherche Médicale (Programme "équipe FRM”) and the LABEX LIFESENSES (reference ANR-10-LABX-65) supported by French state funds managed by the Agence National pour la Recherche within the Investissements d'Avenir programme under reference ANR-11-IDEX-0004-02. F P-B thanks and dedicates this work to Mrs. María González-Mundi.

## Author Contributions

F P-B, YZ, DKS, IAG, AC and KJM designed experiments and analysed results and wrote the paper. F P-B, YZ, DKS and IAG carried out experiments.

## Supplemental Figures

**Supplementary Figure 1.**
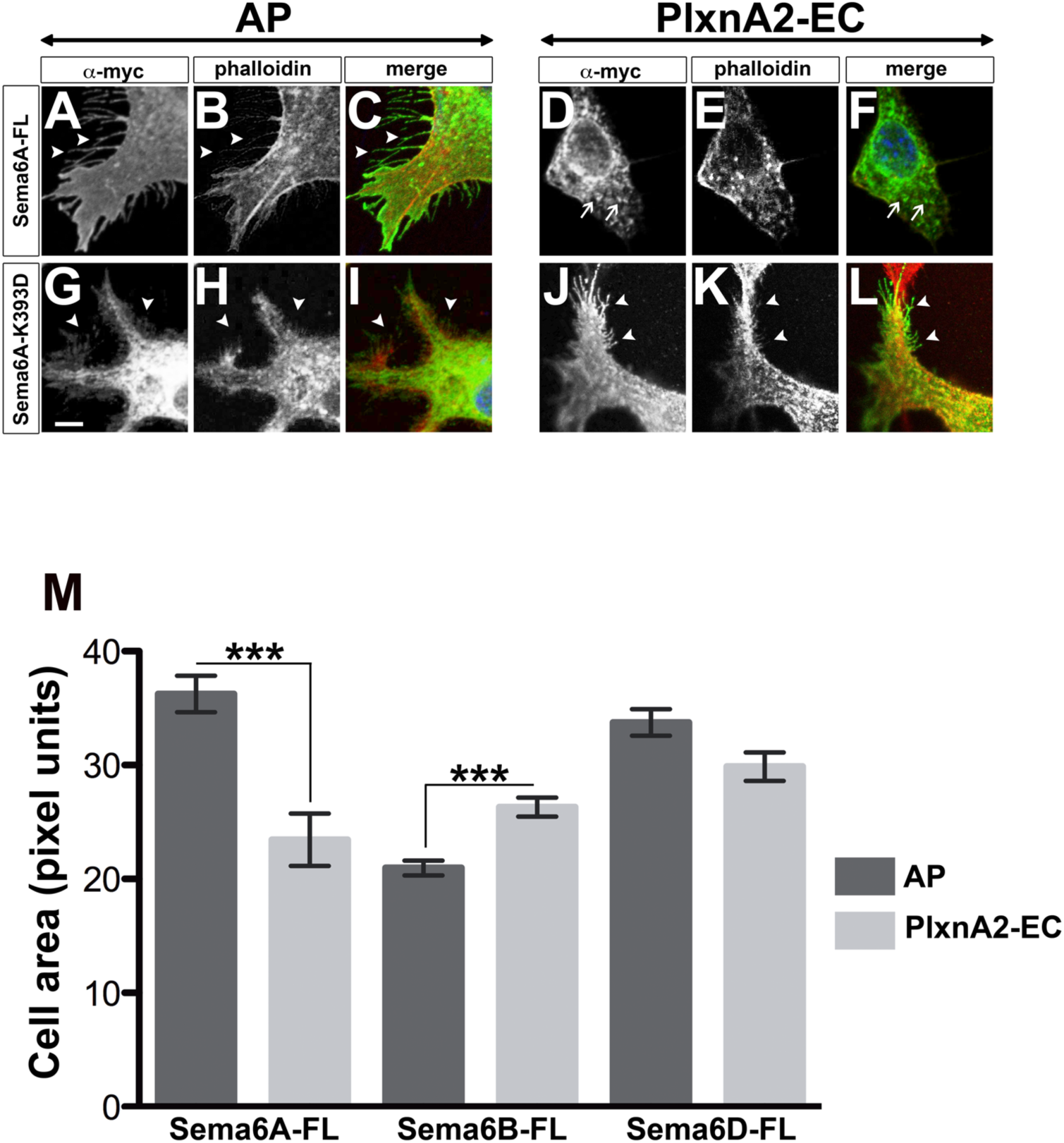
PlxnA2-Sema6A interaction causes cell contraction and actin collapse. (**a-l**) NIH3T3 cells expressing myc-Sema6A-FL (Sema6A-FL; **a-f)**, or myc-Sema6A-K393D (Sema6A-K393D; **g-l**) were treated with purified AP-Fc (AP; **a-c** and **g-l**) or PlxnA2-EC-Fc (PlxnA2-EC; **d-f** and **j-l**) and stained with α-myc antibody and phalloidin to visualise actin fibres. Arrowheads indicate membrane protrusions, and arrows point to collapsed actin clumps. Scale bar = 5 μm. (**m**) Graph represents the cell area in NIH3T3 transfected with Sema6A-FL, Sema6B-FL or Sema6D-FL and treated with AP or PlxnA2-EC; *n*=100–200 cells per experimental condition. Data are expressed as mean ± s.e.m; \*\*\**P* < 0.001; Student’s *t*-test.

**Supplementary Figure 2.**
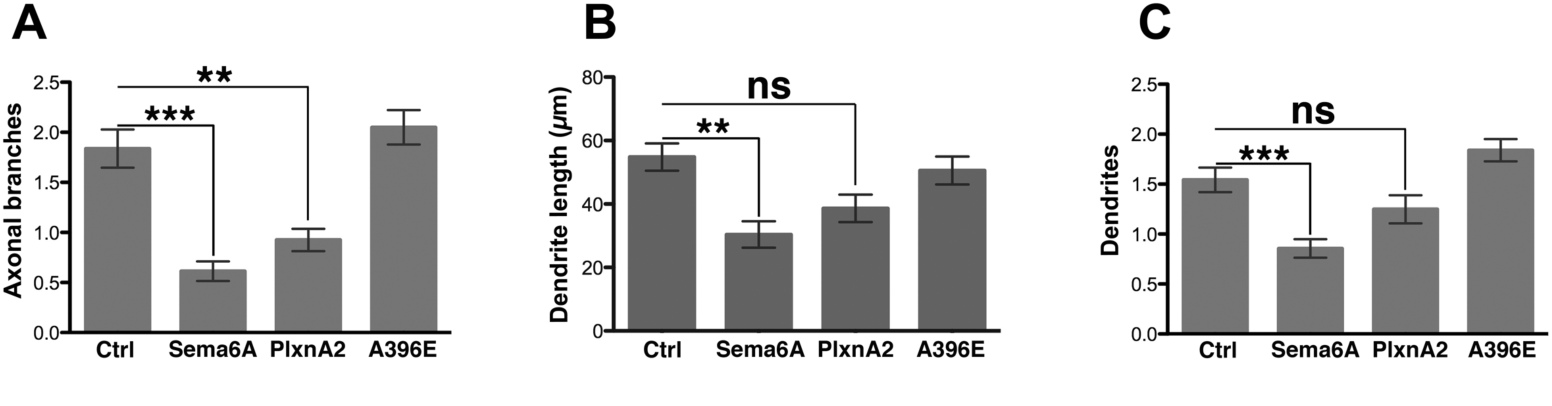
Growth on Sema6A or PlxnA2-expressing substrate reduces the complexity of axonal processes of cerebellar granule neurons. **(a)** Graph represents the number of axonal branches of granular neurons cultured on different NIH3T3 layers. **(b)** Graph represents the dendrite length of granular neurons cultured on different NIH3T3 layers. (**c**) Graph represents the number of dendrites per neuron in granular neurons cultured on different NIH3T3 layers; *n*=100–200 cells per experimental condition. Data are expressed as mean ± s.e.m; \*\**P* ≤ 0.005, \*\*\**P* < 0.001 an ns = *P* > 0.05; one-way ANOVA followed by Bonferroni multiple comparison test.

**Supplementary Figure 3.**
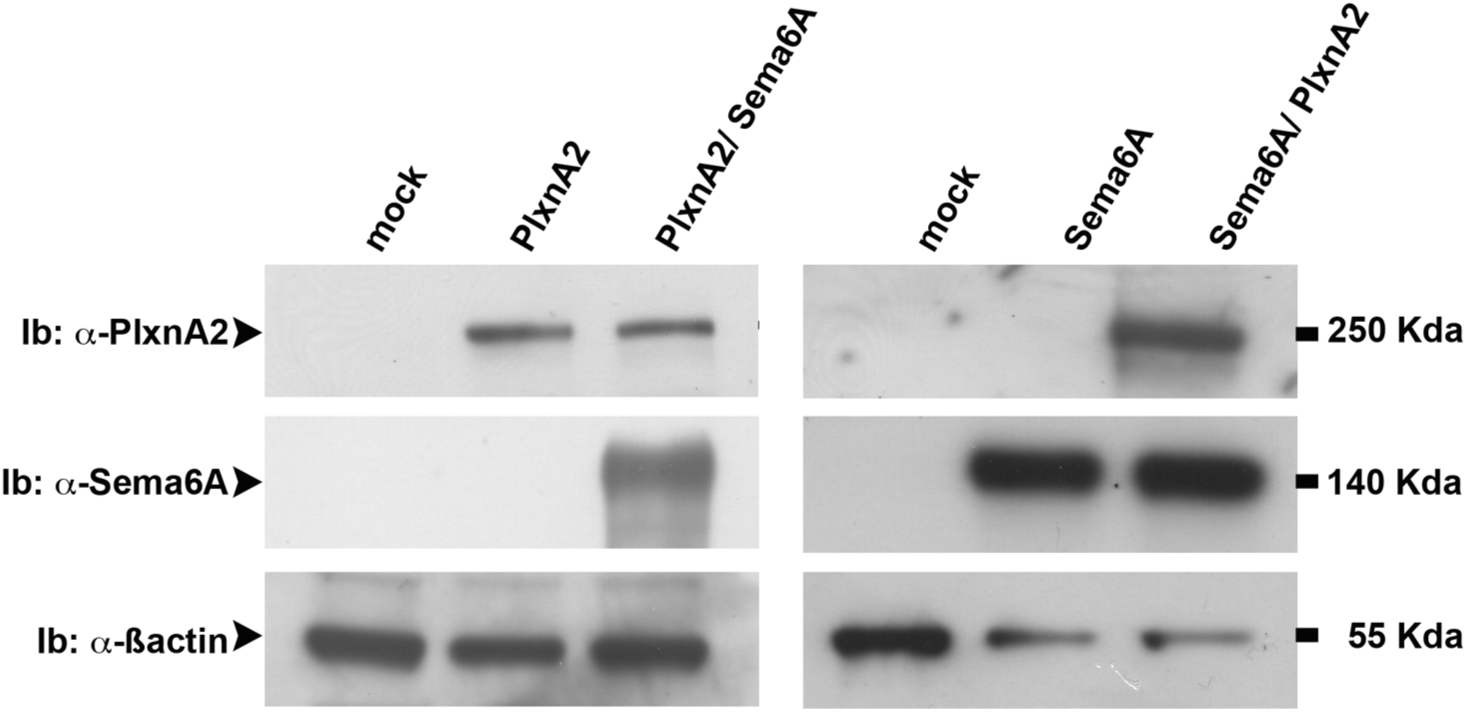
Co-expression of PlxnA2 or Sema6A does not affect expression levels. Immunoblots of total lysates from transfected COS-7 cells expressing PlxnA2, Sema6A or PlxnA2 and Sema6A shows roughly comparable levels of expression of the two proteins and no change in expression levels when they are co-expressed.

**Supplementary Figure 4.**
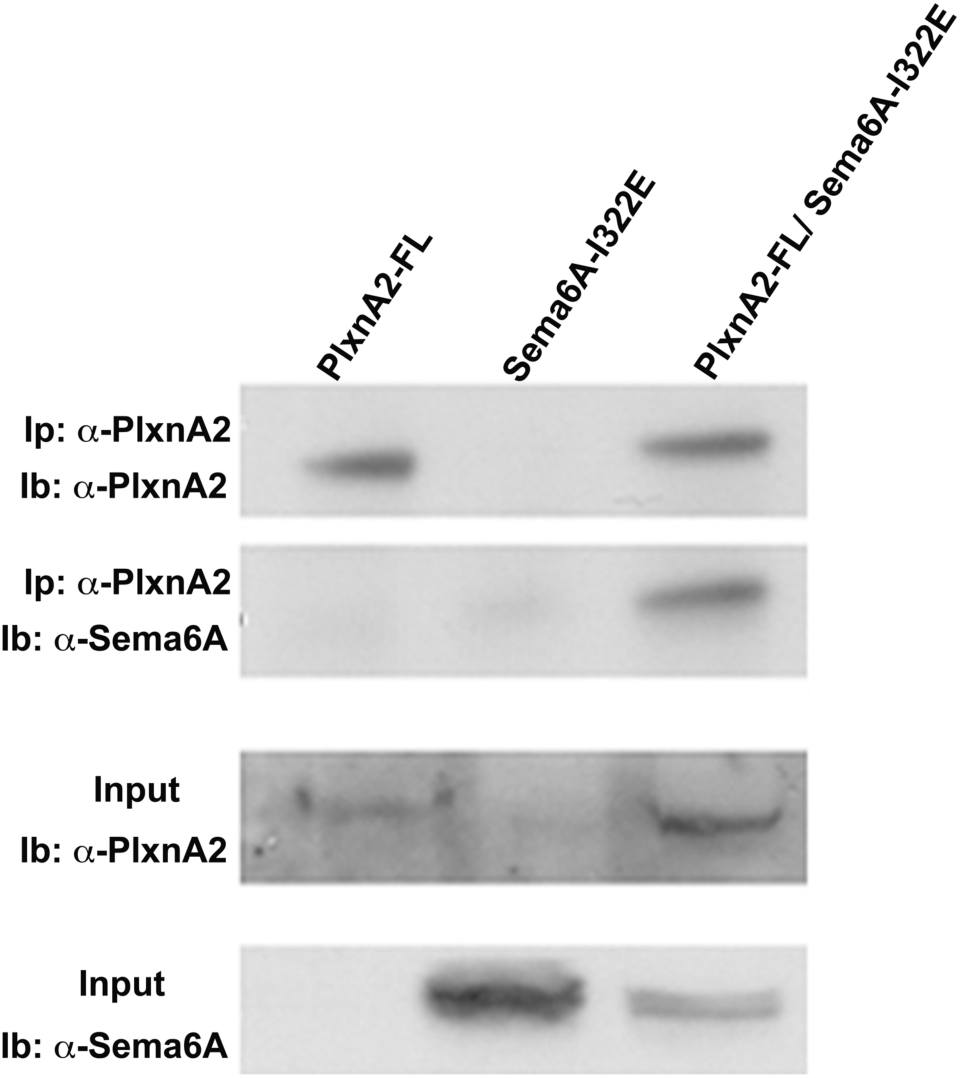
Sema6A monomers can still interact with PlxnA2 in *cis*. PlxnA2 Immunoprecipitations (Ip) from protein samples of COS-7 cells transfected with PlxnA2-FL, Sema6A-I322E or PlxnA2-FL together with Sema6A-I322E. Antibodies against PlxnA2 and Sema6A were used in the immunoblots (lb).

**Supplementary Figure 5.**
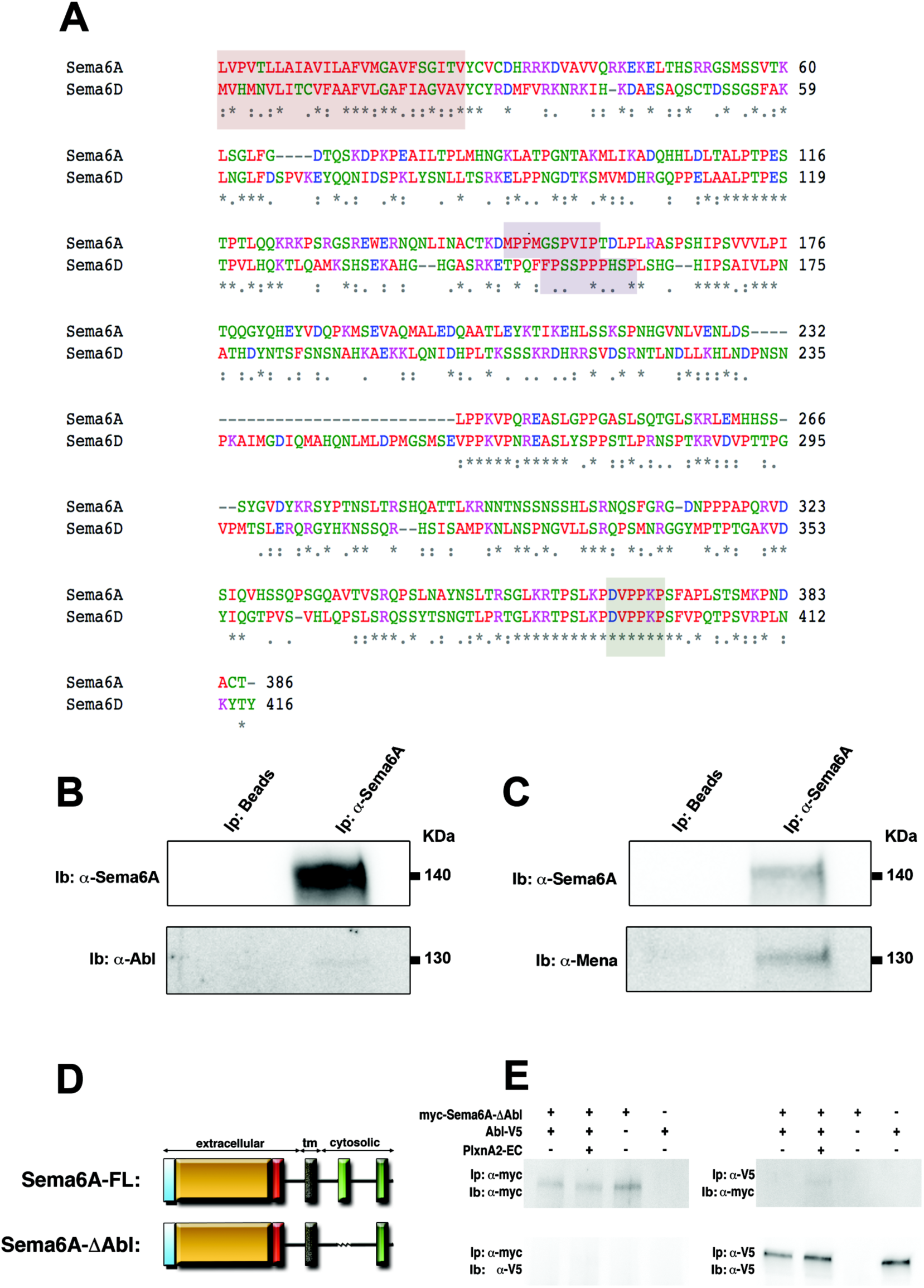
The cytosolic sequence of Sema6A contains Abl and Evl-binding domains. (**a**) Alignment of the cytosolic domains of Sema6A and Sema6D. Orange outline = transmembrane domain; purple outline = Abl binding domain; green outline = Evl binding domain. (**b**) Abl protein is co-immunoprecipitated with anti-Sema6A antibodies from extracts of P6 cerebellum (top); control blot with anti-Sema6A (bottom). (**c**) Mena protein (at ~95kDa) is also co-immunoprecipitated with anti-Sema6A antibodies from extracts of P6 cerebellum (top); control blot with anti-Sema6A (bottom). (**d**) Scheme indicates the structure of Sema6A-FL and Sema6A-ΔAbl. (**e**) Immunoprecipitations from untreated or PlxnA2-EC-treated COS-7 cells transfected with different combinations of myc-Sema6A, Abl-V5. Immunoprecipitations (Ip) and Immunoblots (Ib) were performed employing α-myc and α-V5 antibodies.

**Supplementary Figure 6.**
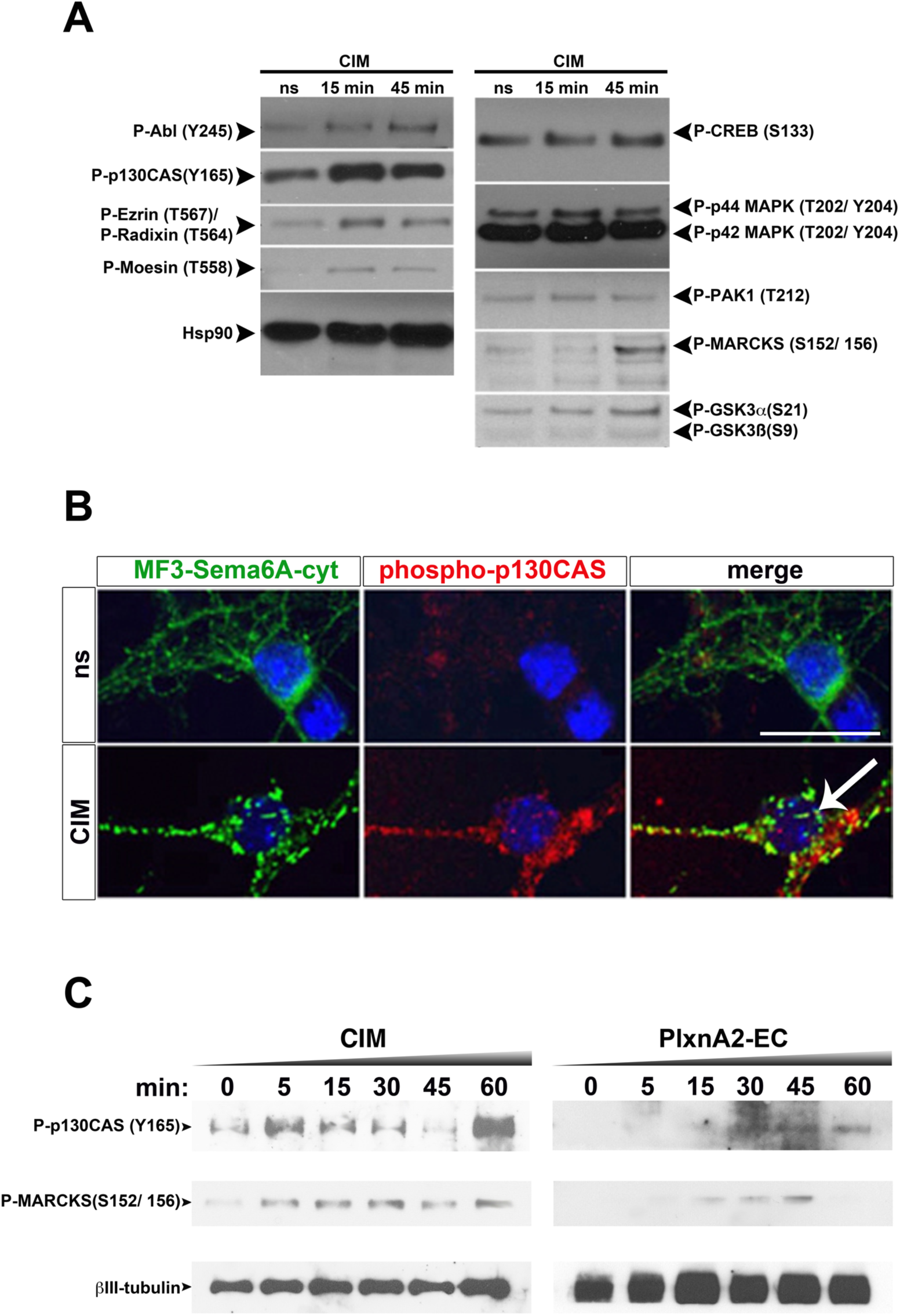
Sema6A multimerisation stimulates the phosphorylation of cytoskeletal interacting proteins. (**a**) Representative immunoblots from MF3-Sema6A-cyt-expressing cerebellar neurons treated with CIM for different lengths of time (min). The phosphorylation of diverse signaling cues was tested by immunoblotting. Hsp90 was used as loading control. (**b**) The clustering of Sema6A-cyt induces the phosphorylation of p130CAS. Moreover, Sema6Acyt clusters co-localised with α-P-p130Cas signal (arrow). Scale bar = 20 μm. (**c**) Immunoblots from MF3-Sema6A-cyt-expressing cerebellar neurons treated with CIM, and cerebellar neurons treated with PlxnA2-EC for different lengths of time (min). The phosphorylation of p130CAS and MARCKS were evaluated with specific phospho-antibodies. βIII-tubulin was used as loading control.

